# Fine-Scale Models of Bee Species Diversity and Habitat in New York State

**DOI:** 10.1101/2024.07.15.603570

**Authors:** Mark A. Buckner, Erin L. White, Timothy G. Howard, Matthew D. Schlesinger, Bryan N. Danforth

## Abstract

Anthropogenic drivers of global change threaten bee diversity and the ecosystem services bees provide. Despite their importance, the conservation of bee pollinators is complicated by limited and often heavily biased occurrence data. A recent state-wide survey of insect pollinators across New York, United States generated a large spatial dataset of bee species occurrence records from community scientists, historical collections, and survey efforts. Using a combination of the state survey records with occurrence data from across the contiguous United States, we applied an ensemble modeling approach using balanced random forest and small bivariate generalized linear models to predict the distributions of most of the state’s bee species. We predicted the spatial distribution of bee species richness using a stacked species distribution model with climate, land cover, and soil covariates. To inform bee diversity conservation, we predicted spatial variation for each species and groups of species sharing similar life history traits. We also estimated statewide distribution of range-size rarity, ecological uniqueness, and climate exposure. We found that the richness of modeled species is high across the state, with the greatest richness in regions with low soil clay content and intermediate forest cover. The fine spatial scale and extent of our gridded data layers match the scale of conservation action in the state, providing an opportunity to incorporate wild bee diversity into broader statewide conservation planning. Conserving New York State’s bee pollinators is not straightforward, and decisions should be based on broader conservation priorities that incorporate bee biodiversity indicators into decision-making. Here, we encourage the inclusion of these vital pollinators in conservation decisions by leveraging the best available data and methods robust to small sample sizes to provide spatially explicit data products representing the distribution of bee diversity across the state of New York.

## Introduction

The broad-scale loss of biodiversity across taxonomic groups is a global phenomenon of significant human and ecological concern (Sage, 2020). The threat of biodiversity collapse has highlighted the need to protect economically and ecologically important species including vital insect pollinators such as bees (Hall & Steiner, 2019). Public and political concern regarding bee decline came to the forefront with the onset of colony collapse disorder in the mid-2000s (Althaus et al., 2021; vanEngelsdorp et al., 2009). Consequently, pollinator protection initiatives in the United States have generally focused on *Apis mellifera*, the managed western honeybee (Colla & MacIvor, 2017; Iwasaki & Hogendoorn, 2021). While such initiatives have bolstered public interest in addressing pollinator declines, misguided or even counterproductive calls for honeybee conservation disregard the loss of native bees (Geldmann & González-Varo, 2018). Yet, our awareness of humanity’s role in unmanaged wild bee declines is not new (Carson, 1962; Frison, 1926; Williams, 1982), nor is our understanding of the consequences of losing wild pollinators, with the loss of wild bee mediated pollination services noted since at least the mid-1900s (Carson, 1962; Fronk & Painter, 1960; Stephen, 1955). To date, pesticide use, pests and pathogens, habitat loss, introduced species, and climate change, among others, have all been implicated in global bee diversity declines (Dicks et al., 2021; Goulson et al., 2015; Potts et al., 2010; Vanbergen et al., 2013).

Considering the growing need to address bee declines, national or regional survey efforts may prove vital for establishing conservation baselines, formulating priorities, and enabling status assessments (Woodard et al., 2020). Although the utility of survey efforts may be limited in the near term (see Tepedino & Portman, 2021), they present a clear opportunity to address historical data limitations that have hindered native bee conservation efforts (Inouye et al., 2017). While global analyses of bee diversity exist (Bystriakova et al., 2018; Orr et al., 2021), accounting for fine-scale, local variations through national or subnational assessments could better inform conservation action (Schmidt-Traub, 2021). For example, there is growing evidence of the importance of forests for pollinator diversity (Ulyshen et al., 2023), despite global bee diversity hotspots primarily occurring in midlatitude arid deserts (Bystriakova et al., 2018; Orr et al., 2021).

In 2021, the New York Natural Heritage Program concluded the Empire State Native Pollinator Survey (ESNPS), a four-year statewide survey focused on evaluating the conservation status of New York State’s (NYS) insect pollinators. The effort resulted in the aggregation of numerous biodiversity datasets, community science contributions, and surveys to determine the status of NYS bee, fly, beetle, and moth pollinators. To evaluate conservation statuses, the survey relied on coarse distribution trends, rarity, and expert assessment of the threats and vulnerabilities each species faces (Schlesinger et al., 2023; White et al., 2022). The effort highlighted the need for the inclusion of insect pollinators in statewide conservation planning, with 60% of native insect pollinator species ranked as vulnerable or imperiled (White et al., 2022).

State surveys like the ESNPS provide significant contemporary, spatially accurate records of species localities which, when paired with modern species distribution models (SDM), provide opportunities for mapping local bee diversity. Here we integrated the ESNPS bee locality dataset with collection records from across the Contiguous United States (CONUS) to model the distribution of over half of the state’s bee species. We employed ensemble models, with balanced random forest and small bivariate generalized linear model constituents, to generate predicted distributions robust to the small sample sizes and presence-only data available for NYS’s bee pollinators. We predicted the distribution of species richness across the state and identified regions that may be of high conservation value. Regions of high conservation value include those that are important for range-restricted species and represent unique community compositions. In addition, we evaluated the threats posed by climate change to identify the bee communities experiencing the greatest pressure for climate-associated redistribution. Our approach provides a roadmap for augmenting conservation assessments following state-wide survey efforts and presents gridded data products to facilitate the consideration of wild bee pollinators in conservation planning.

## Materials and Methods

### Bee Occurrence Data

White et al. (2022) and Schlesinger et al. (2023) presented a thorough overview of the data collection and integration process used in the ESNPS dataset which consists of records from curated datasets (Richardson, 2024), natural history museums, community science observations (i.e., iNaturalist), and structured surveys across the state’s ecoregions. To better represent the full distribution of environmental conditions NYS bee species occupy, we supplemented the ESNPS dataset with records from across CONUS obtained from the Global Biodiversity Information Facility (GBIF; GBIF.org, 2024). We performed all analyses in R v4.1.2 (R Core Team, 2021).

We checked for georeferencing errors with COORDINATECLEANER v3.0.1 (Zizka et al., 2019) by flagging coordinates that correspond to country or state centroids and biodiversity institutions. We chose to retain records from botanical gardens, museums, and university campuses as we found that the audited records corresponded to collecting events occurring on institution campuses. In addition, we removed any records within 100m of county centroids unless they included additional locality information (e.g., ‘Beebe Lake, Tompkins County, NY’). We chose to retain records for species that have not authoritatively been identified in NYS as these may constitute early records or range expansions. Finally, we filtered the dataset to retain observations between 1971 and 2023 for species that, after removing duplicate observations, had a minimum of ten spatially unique records.

### Environmental Covariates

We considered fourteen environmental covariates that are likely to influence bee species distributions (Table S1). To ensure the greatest possible temporal overlap between the climate data and observation records, we selected climate covariates derived from the 1981-2010 climate normals available in CHELSA v2.1 (Karger et al., 2017, 2021). Besides climate, we included percent canopy cover (LANDFIRE, 2014) and soil texture data (Hengl, 2018a, 2018b) and derived two additional layers representing gridded terrain roughness as the standard deviation of the slope (Grohmann et al., 2011; LANDFIRE, 2020) and aridity index calculated as the ratio of annual precipitation to potential evapotranspiration (United Nations Environment Programme, 1992).

The optimal grain for SDMs should reflect the ecological scale at which the species of interest respond to their environment, even if this is less than the spatial uncertainty of the occurrence data (Gábor et al., 2022). The scale at which bees interact with their environment corresponds closely with body size and sociality (Greenleaf et al., 2007; Kendall et al., 2022). Most bees forage within less than 1 km of their nest, with large, social species often foraging between 1-2 km (Kendall et al., 2022). Since bee records often represent foraging individuals, we accounted for the environmental conditions controlling foraging and nesting habitat by aggregating each gridded dataset to approximately 990-m resolution and projected the data layers to an Albers equal area projection.

To ensure that records from wetlands were not excluded erroneously for occurring within waterbodies, we categorized cells as water only if more than 90% of the aggregated cells in a 30-m resolution vegetation layer were classified as water (LANDFIRE, 2014). Where required, we imputed the soil data surrounding water bodies using the mean of a three-by-three focal window in TERRA v1.7-65 (Hijmans, 2023). Finally, we conducted principal components analysis (PCA) to mitigate the effects of collinearity and reduce the dimensionality of the environmental variables. To limit the number of constituent models, thereby reducing computational cost, we included the minimum number of components required to explain 80% of the variance in the dataset.

### Background Points and Bias Correction

We used two bias correction methods to address spatial sampling bias in the opportunistic collection records. First, we calculated a bias layer with kernel density estimation using a 20-km bandwidth across CONUS. From this bias layer, we selected 50,000 weighted random background points with similar distribution and spatial bias to the presence records (Inman et al., 2021; Kujala et al., 2015; Valavi et al., 2021). Due to the variability of sampling effort, we used an additional model-based bias correction step where we modeled sampling intensity with a spatial nonstationary Poisson point process model in SPATSTAT v3.0-6 (Baddeley et al., 2015). Similar model-based methods have been shown to improve predictive performance and have been widely used outside of distribution modeling (Warton et al., 2013). We chose covariates that represent where researchers and community scientists often search for bees, including population density (Center for International Earth Science Information Network - CIESIN - Columbia University, 2018), protected area status (USGS GAP, 2022), distance to primary and secondary roadways (U.S. Census Bureau, 2013), distance to biodiversity institutions (Zizka et al., 2019), and whether the grid cell is part of the Fort Drum military base (U.S. Census Bureau, 2010)— a site with additional high intensity sampling during the ESNPS. In addition, we included polynomial terms for the spatial coordinates and interactions between the distance variables and protected area status. We included the gridded intensity layer as a covariate in the SDMs. When predicting across space, we set the intensity layer equal to its global mean such that distribution predictions assume all cells had identical sampling intensit

### Stacked Species Distribution Models

Due to the small number of records available for many bee species, following conventional recommendations to avoid overfitting (e.g., a minimum of 10 presence-only records per covariate) would necessitate fitting unrealistically simple models (Brun et al., 2020). To overcome this limitation, we integrated additional records from across CONUS and trained separate ensemble models for each species. Each ensemble consisted of a balanced random forest (BRF) model and a collection of small bivariate generalized linear models, referred to as an Ensemble of Small Models (ESM). Both constituent model types, and ensemble models in general, have shown promise for modeling rare species (Breiner et al., 2015; Erickson & Smith, 2023; Lomba et al., 2010; Mi et al., 2017; Valavi et al., 2022). Additionally, ensemble models have been found to perform adequately with presence-only datasets, with BRF and regression methods both performing well with small sample sizes and comparably well to other model types when large numbers of presence points are available (Valavi et al., 2022).

For each species, we trained the BRF models as probability forests in RANGER v0.15.1 with 1,000 trees and all other parameters set to their defaults (Wright & Ziegler, 2017). By accounting for class imbalance during training, BRFs perform better than conventional random forest models when fitting SDMs with small numbers of presence localities (Valavi et al., 2022). Additionally, we fit the ESMs as downweighed bivariate generalized linear models representing the pairwise combinations of all covariates for each species. We trained two sets of ESMs, one with linear terms only and one with both linear and polynomial terms for each pairwise combination of covariates. While including polynomial terms increases the computational cost of model fitting significantly, they can perform better than linear-only ESMs for discrimination tasks (Erickson & Smith, 2023).

We estimated ensemble model performance using the cross validated area under the receiver operating characteristic curve (AUC_ROC_) for each model. For species with greater than 50 records, we used 5-fold spatial block cross-validation, and for species with fewer than 50 records, we used nearest neighbor distance matching, a modified version of leave-one-out cross-validation (Milà et al., 2022; Valavi et al., 2019). Occasionally spatial cross-validation returned invalid folds, where one or more folds contained only a single test point. In such cases, we applied buffered leave-one-out cross-validation or, if all spatial methods failed to return valid folds, stratified k-folds implemented with CARET v6.0-94 (Kuhn, 2008). All spatial cross-validation methods were performed with BLOCKCV v3.1-3 (Valavi et al., 2019). We generated folds for each species independently and used identical folds across all constituent models.

To assemble the final ensembled predictions, we took the weighted average of each constituent model with Somers’ D where *D* = 2 ∗ (*AUC* − 0.5), and excluded models where *D < 0* (Breiner et al., 2015). We calculated the final cross validated AUC_ROC_ for each species based on the same weighting scheme as the ensemble predictions. We then stacked the predicted distributions of each species by taking the sum of the occurrence probabilities at each grid cell (Calabrese et al., 2014). Approximating species richness in this way allowed us to estimate the variance in site-level richness by assuming richness follows a Poisson Binomial distribution; however, we were unable to include uncertainty from the occurrence probabilities themselves which are assumed fixed in this stacking scheme (Calabrese et al., 2014).

### Other Biodiversity Indicators

We identified areas that may be of high conservation value for NYS bees based on species richness, ecological uniqueness (the contribution of cell-level community composition to overall beta diversity), and range-size rarity (RSR; the importance of each grid cell for range-restricted species). Where possible, we computed these metrics using probabilistic distribution predictions instead of binary threshold predictions, as using a threshold does not improve occupancy estimates and may lead to over-prediction if performed before stacking (Calabrese et al., 2014; Guillera-Arroita et al., 2015). However, to estimate community composition, we applied probability ranking rules to threshold the probability predictions, resulting in presence-absence maps for each species. Probability ranking rules are a community-based method to threshold SDM predictions where species are marked present in each grid cell up to the predicted species richness based on their relative occurrence probabilities within each grid cell (D’Amen et al., 2015).

We calculated ecological uniqueness in JULIA v1.9.4 as the cell-wise local component of beta diversity, where more unique community compositions were found in cells with greater relative contribution to statewide beta diversity (Dansereau et al., 2022; Legendre & De Cáceres, 2013). In addition, we calculated RSR to identify cell-level importance for range-restricted species (Guerin & Lowe, 2015). We calculated RSR using the continuous occurrence probability predictions, by first estimating each species’ occurrence area as the sum of occurrence probabilities across CONUS (Stark & Fridley, 2022). Then, to estimate the cell-wise importance of each grid cell to range-restricted species, we took the sum of each species’ occurrence probability divided by their occurrence area.

To assess the potential effects of climate change, we calculated the distance-based velocity of climate change (VoCC) between the 1981-2010 and 2071-2100 climate normals across NYS as the mean of four General Circulation Models (GFDL-ESM4, MPI-ESM1-2-HR, IPSL-CM6A-LR, MRI-ESM2-0) for two shared socioeconomic pathways representing the most optimistic (SSP1-2.6) and pessimistic (SSP5-8.5) scenarios from Coupled Model Intercomparison Project Phase 6 (CMIP6). We applied a 250-km buffer zone around NYS to ensure that analogous climates were identifiable within a reasonable dispersal range. We used multivariate VoCC based on three climate variables: mean annual air temperature, annual precipitation, and growing season precipitation following the multivariate method described by Hamann et. al (2015). To calculate species-specific climate exposures, we used the mean cell-wise climate velocity across NYS weighted by occurrence probability for each species. Next, we identified the regions of the state that have the greatest community-wide risk from climate change as equation 1:

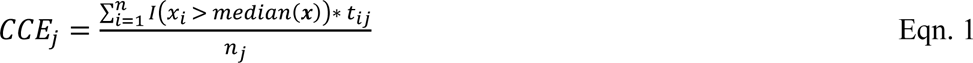

where community climate exposure (*CCE*) at the *j*-th site is equal to the proportion of the total species (*n*), which are present (*t*_*ij*_ = 1) in the threshold predictions (**T**) that have a ratio of climate exposure to range size (***x***) above the median ratio across all species.

To evaluate the utility of permanently protected areas for bee conservation, we assessed how bee species richness, RSR, ecological uniqueness, and climate exposure within protected areas deviated from the statewide mean. The US Geological Survey protected area inventory assigns protected areas a status code depending on the permanency of protection and allowable uses (USGS GAP, 2022). GAP status 1 and 2 are under mandates to ensure permanent protections and differ from GAP status 3 areas that are permanently protected but allow extractive uses (e.g., logging or mining) (USGS GAP, 2022). For this analysis, we focused on protected areas with GAP statuses 1, 2, and 3 as delineated by the New York Protected Area Database (New York Natural Heritage Program, 2024).

## Results

Our complete bee occurrence dataset contained over 213,000 unique records representing 269 bee species. The number of presence-only records varied between a minimum of eleven for *Osmia felti* and over 35,000 records for the common eastern bumble bee, *Bombus impatiens*. Approximately 21% of the included species had 50 or fewer unique records across CONUS during the 52-year data period (1971-2023) and 79% of all observations were recorded after 2010. The occurrence records showed clustering in and around urban areas and roadways suggesting significant spatial sampling bias (Figure S1). While many species had insufficient data to include in the models, we noted a similar distribution of species’ life history traits in the modeled species as compared to all known species from NYS, although solitary bees were overrepresented in our dataset (Figure S2).

Following dimensionality reduction with a PCA, the first four principal components accounted for 80.6% of the total variance in our environmental layers (Table S2). Principal Component (PC) 1 primarily measured temperature, showing strong positive associations with climate variables representing annual and winter temperatures and a negative association with annual temperature range. PC2 showed positive associations with aridity index and percent canopy cover. PC3 primarily represented soil texture and PC4 was dominated by growing season precipitation. Predicted sampling intensity, from the point process model, was highest near roadways and biodiversity institutions with notable peaks in intensity along roadways near institutions, in Fort Drum, and near New York City (Figure S3).

We fit two separate ensemble models which varied in whether the ESM constituents consisted of linear terms only or linear and polynomial terms. We used spatial cross-validation to evaluate model performance for 97.8% of the included species. The linear-only models performed slightly worse on average (mean AUC_ROC_: 0.72; range: 0.51-0.96) than the models that included linear and polynomial terms (mean AUC_ROC_: 0.74; range: 0.50-0.96). Models of target species during the ESNPS performed better on average (AUC_ROC_: 0.75) than non-target species (AUC_ROC_: 0.72). We were unable to fit satisfactory models for two species, *Andrena heraclei* and *Epeolus autumnalis,* when using linear terms only (i.e., all constituent models had an AUC_ROC_ < 0.5), and all models failed to reach a satisfactory AUC_ROC_ for two additional species, *Apis mellifera* and *Macropis nuda*. Models of *Protandrena andrenoides* were fit without all available GBIF records due to an unidentified taxonomic conflict between the datasets. Species that have not previously been authoritatively identified within NYS (n = 8) showed low predicted suitability within the state.

The predicted richness of the modeled bee species across the state was relatively homogeneous, varying between 75 and 150 species (Interquartile range: 13.3 species) with a left-skewed distribution (Figure 1a). Bee richness was high throughout the state, peaking in the Adirondack foothills of northern NYS. In the remainder of the state, richness was moderately positively correlated with percent canopy cover (r = 0.67) and negatively correlated with soil clay content (r = -0.75). Correspondingly, we found that species richness peaked at intermediate values of PC3, dominantly representing soil texture, and PC2, representing canopy cover and aridity (Figure S4). Our models predict low richness in the heavily agricultural Finger Lakes Region, with the lowest predicted richness occurring in the clay rich lowlands west of the Adirondacks and the “Black Dirt” region near the Hudson Valley in southern NYS. Long Island had moderately low predicted richness compared to nearby inland areas such as the Hudson Valley Ecoregion and surrounding foothills. The variance in the stacked models was even across the state with minimal variation in uncertainty from the stacking process (Figure 1b). We subset the overall richness predictions into subgroups by taxonomy or life history, including *Bombus* spp., host plant specialization, sociality, native and non-native species, nesting behavior, and apple pollinators. Except for non-native (r = -0.09), eusocial (r = 0.63), and *Bombus* species (r = 0.69), the spatial distribution of each subset showed a strong positive correlation with the prediction for all species (r > 0.95).

**Figure 1:**
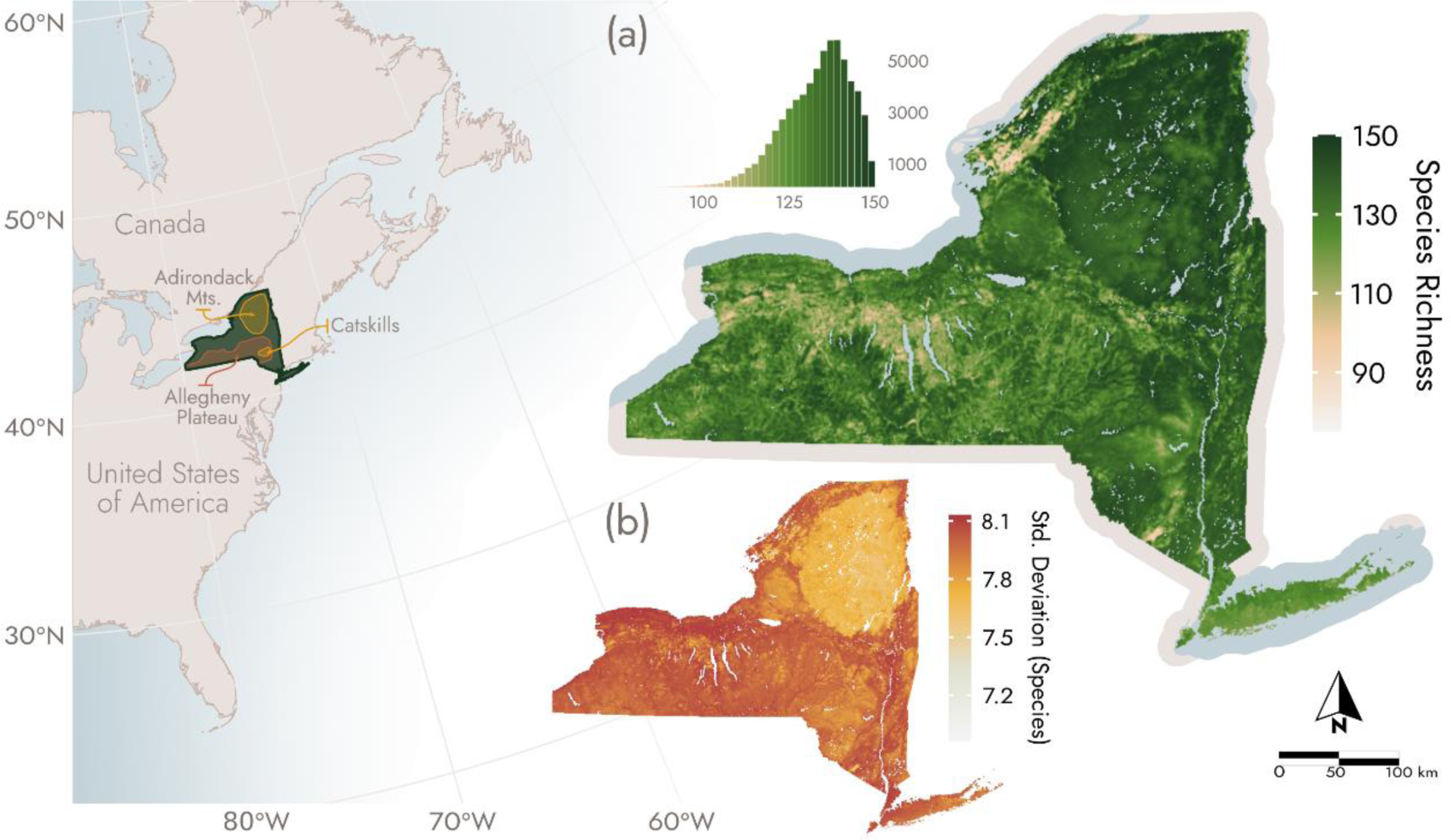
(a) Predicted richness is relatively high across much of the state with low species richness in agricultural areas with high clay soil content. The histogram shows the distribution of richness values. (b) Cell-level standard deviations of species richness across the state representing uncertainty from the stacking process.

In addition to bee richness, we evaluated the spatial variation in the importance of each grid cell for range-restricted species and ecological uniqueness, calculated as the local component of beta diversity. We found that the spatial distribution of RSR was similar to species richness (Figure 2a). In contrast, we found that community compositions were most unique in regions with low richness, including the Finger Lakes and coastal Long Island (Figure 2b). Uniqueness was also higher in some urban areas, including Albany, New York, though interestingly it was not comparably high in the ecologically unique Albany Pine Bush Preserve. Overall, all bee communities showed a strong overlap with 63% of species shared between cells with the most and least unique community compositions.

**Figure 2:**
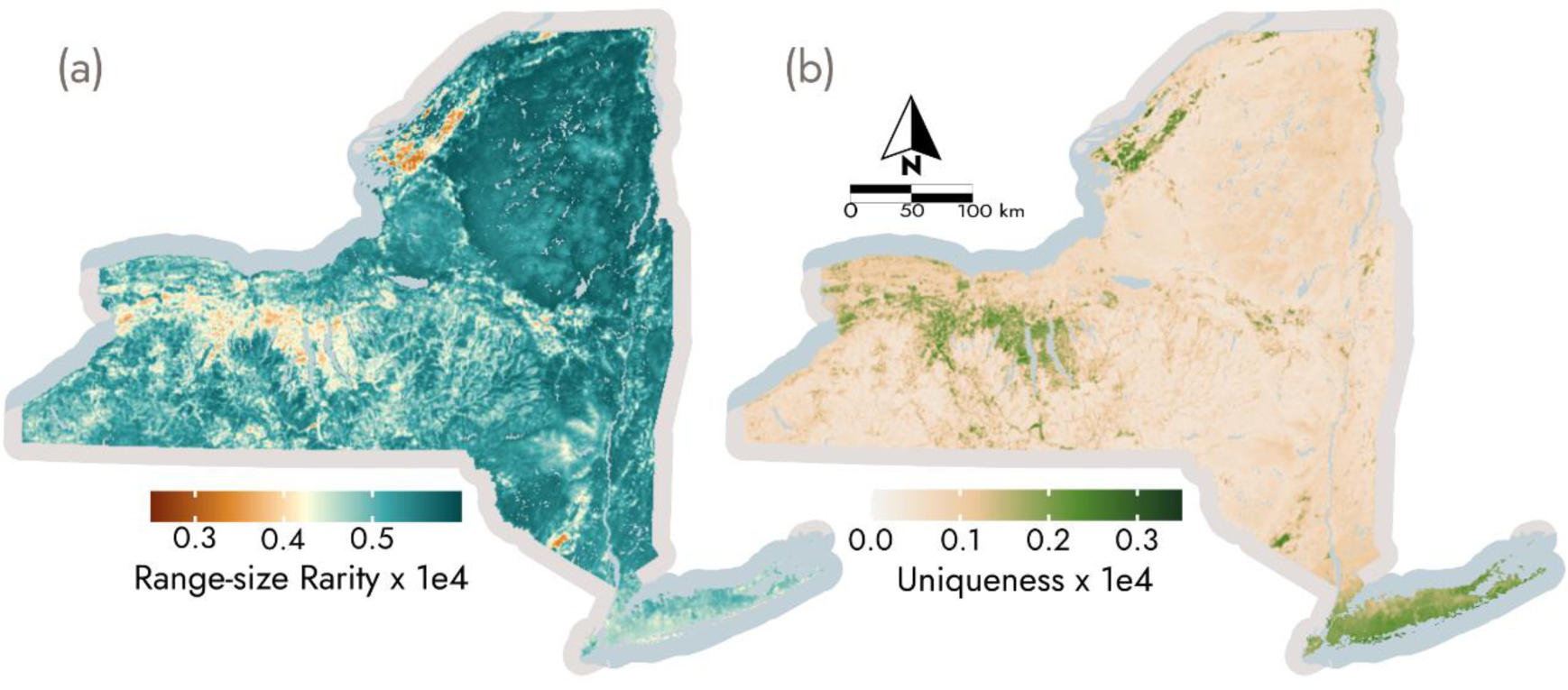
(a) Range-size rarity is lowest in regions with low bee richness. (b) Ecological uniqueness of bee community predictions is higher in areas with low predicted species richness and across Long Island. Both metrics are multiplied by 10,000 for visualization.

Bee communities found at relatively high elevations, particularly within the Adirondacks, Catskills, and Allegheny Plateau, face high climate exposures with lower exposures in the Finger Lakes Region and lowlands south of Lake Ontario (Figure 3b). Overwhelmingly, we expect most bees across the state to track their climate niche northward (Figure 3a). The Adirondack foothills, however, present a clear exception to this pattern where climatic niches displace upward in elevation, even if this means a southward trajectory. This pattern breaks down under the more pessimistic climate change scenario (SSP5-8.5) where 24% of the state, including the Adirondacks, has no climate analog within the 250-km buffer zone, compared to less than 0.1% under SSP1-2.6 (Figure 3c). A sizable portion of NYS’s protected areas are within the Adirondacks, contributing to higher CCEs within protected areas than the statewide average (Figure 4).

**Figure 3:**
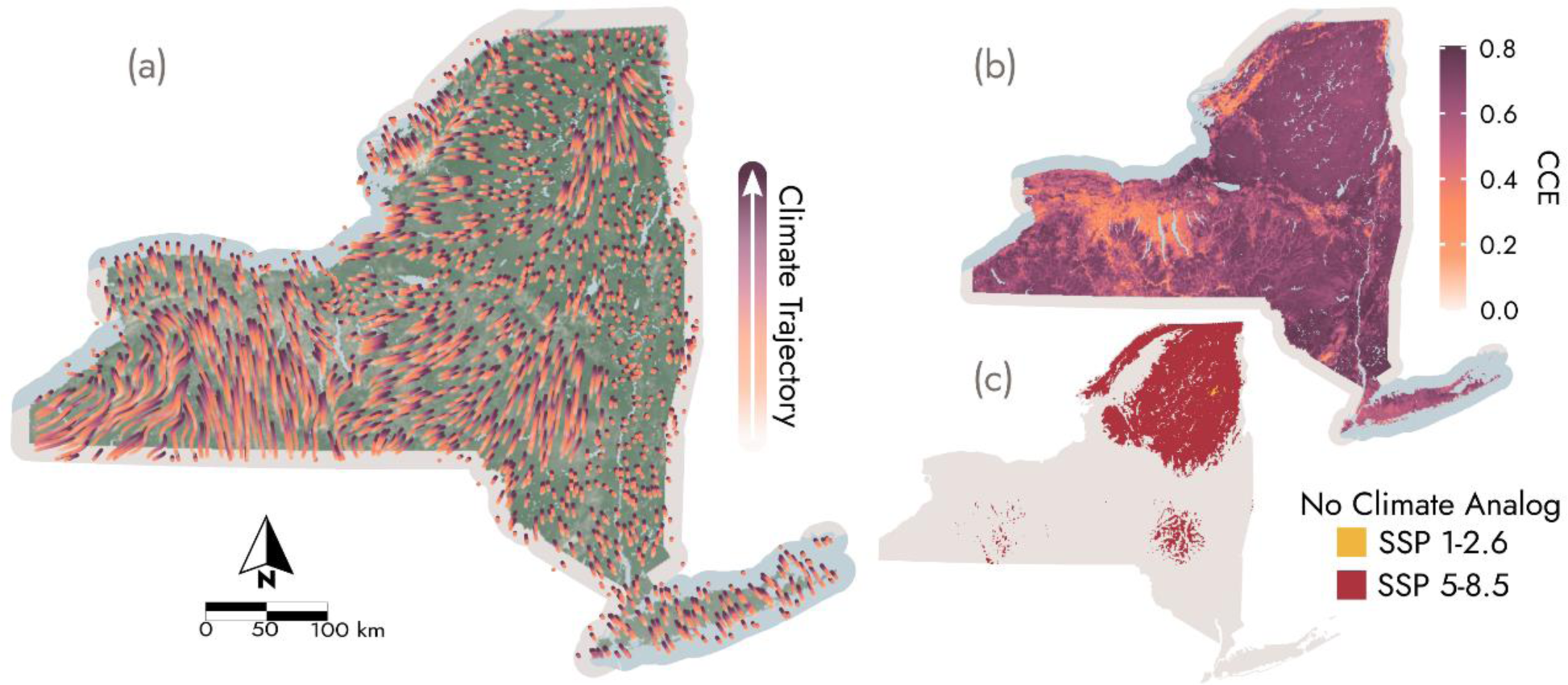
(a) Climate trajectories indicate that climate niches are expected to predominantly move northward under SSP 1-2.6. Streamline length represents the velocity of climatic niche movement. (b) CCE under SSP 5-8.5 is high across the Adirondacks, Allegheny Plateau, and Hudson Valley. (c) The proportion of the state without a climate analog within the 250-km buffer zone increases drastically with more extreme climate change.

**Figure 4:**
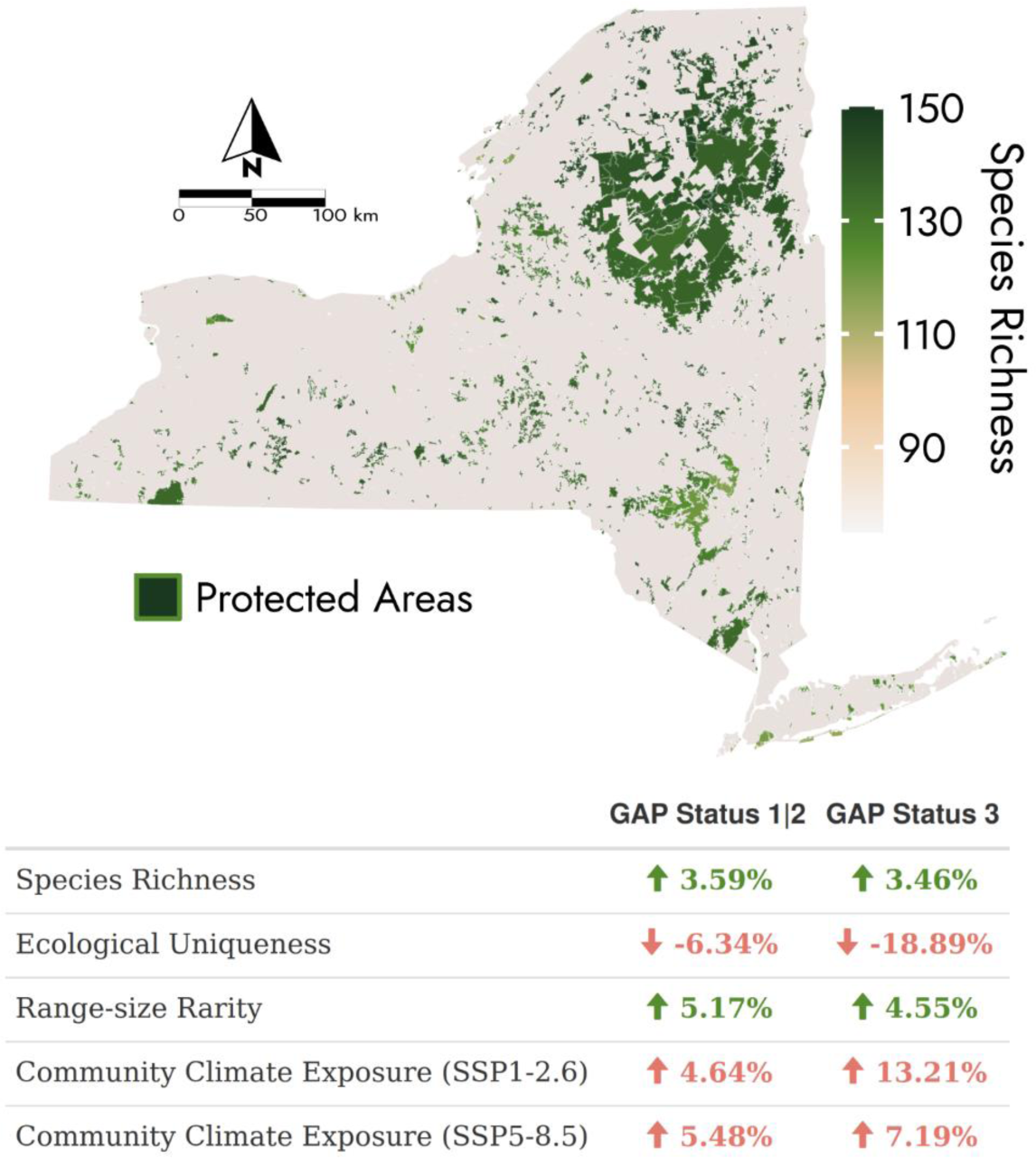
Protected areas across New York State are primarily clustered in the Adirondack Mountains and Catskills. These areas have higher species richness on average than the statewide mean, lower ecological uniqueness, greater importance for range-restricted species, and higher CCEs. The reported percentages are the deviation from the statewide average within protected areas. Areas where extractive uses are permitted (GAP status 3) showed greater deviations from the statewide average in ecological uniqueness and climate exposure than areas prohibiting extraction (GAP status 1 and 2).

## Discussion

The conservation of wild bees faces challenges from deficient occurrence data and limited knowledge about their ecology. Through training SDMs optimized for rare species and robust to spatial sampling bias, we generated fine-scale predictions of bee richness, community uniqueness, climate exposure, and individual species distributions for 60% of NYS’s bee species. The principal results of this study are our model predictions, which we provide as freely available gridded datasets at 990-m resolution (Buckner et al., 2024a). Our model predictions indicated that bee richness was consistently high across NYS and associated negatively with clay soils and positively with percent canopy cover. These patterns are consistent with habitat preference expectations for ground nesting and forest associated species (Antoine & Forrest, 2021; Ulyshen et al., 2023). Conversely, bee communities in regions with high soil clay content and those inhabiting Long Island’s coastal ecosystems were among the most unique communities within the state. Statewide, climate change may contribute to the northward redistribution of bee ranges. A clear exception to this pattern is in the Adirondacks, where climate trajectories point toward movements upward in elevation and no analogous climate within 250 km under SSP5-8.5, indicating a potential climate trap.

The patterns of bee diversity we observed vary from the general expectations presented in global analyses of bee diversity and biogeography, which suggest that bees are most diverse in arid, seasonal habitats (Almeida et al., 2023; Bystriakova et al., 2018; Michener, 1979; Orr et al., 2021). Our findings highlight the importance of modeling biodiversity at scales that permit the inclusion of local ecological contexts. For bees specifically, ensuring effective conservation action requires an understanding of the habitat associations of the regional bee assemblage. In the context of NYS, basing conservation action on the results of global assessments may overestimate the importance of grassland habitats and underestimate that of forests, which our models predict contain considerable bee diversity and which have growing evidence of their role as habitat and forage for spring flying pollinators (Harrison et al., 2018; Ulyshen et al., 2023).

Our work shows the feasibility of developing fine-scale maps for guiding conservation of data-limited taxa like bees, despite insufficient historical survey effort, dependence on unstructured collection records, and taxonomic biases (Chesshire et al., 2023; Rousseau et al., 2024). While monitoring programs are taking shape to address these data deficiencies, many large-scale conservation actions are slated to occur by the end of the decade under the 30x30 framework, that strives for 30% of land area to be conserved by 2030 (Dinerstein et al., 2019). The rapid timeline laid out in these targets presents a challenge for historically underprioritized wild bee species. The spatial data layers we report here are a valuable resource for including native bee diversity in imminent conservation decisions and can inform the protection of these ecologically and economically important pollinators.

In December 2022, the NYS legislature passed a bill modifying the state’s land acquisition policy to align with 30x30 (Thirty by Thirty Conservation Goal, 2022). Given the rapid timeline laid out in this legislation, our results provide an opportunity to include bee diversity as a component of broader conservation planning across the state using the best available data. The current global maps of bee diversity can play a vital role in facilitating cooperative, international conservation prioritizations but are provided at too course of a resolution to inform local conservation action (Chaplin-Kramer et al., 2022; Wyborn & Evans, 2021). Even national biodiversity conservation programs are often hindered by a lack of maps on complementary scales (Schmidt-Traub, 2021). While nationwide maps of bee diversity on similar scales to what we present here are likely feasible, they will require significant data and computational overhead. By focusing on a smaller spatial area, we could achieve fine resolutions of our gridded data layers and therefore provide an opportunity to expand the utility of the recent statewide survey effort.

While the ESNPS provided a source of high-quality, recent bee observations, many of the limitations common to bee occurrence records were still present. Persistent data deficiencies prohibited us from including many of the bee species known from the state. The species that did not meet the minimum requirements are likely rare within NYS or may be specialized on host plants or habitats that are not regularly sampled. As such, we urge caution when considering our predictions of richness and RSR as we were only able to consider the most common and widely distributed species. Similarly, we note that our climate exposure results do not indicate probable species extinctions from an inability to redistribute, as all the modeled species had wide ranges outside of the Adirondacks. To better estimate the threat of climate change, further analyses and data on bee dispersal, adaptive potential, and the influence of biotic interaction are needed.

Presence-only records present a challenge for generating unbiased spatial predictions. By thoroughly accounting for sampling bias in the modeling process, we were able to limit the effects but additional biases from variation in the sampling intensity of different environments may still be present in our models. Moreover, presence-only models are not able to accurately represent demographic rates or abundances (A. Lee-Yaw et al., 2022), imposing limits on the conservation implications of these results. Even so, without implementing large-scale, extensive, and structured repeat surveys, such analyses are not realistic due to the lack of structured historical data. Given the substantial cost and time required to make up for the limitations in existing insect biodiversity data, we emphasize the importance of leveraging the best data currently available, by using methods robust to small sample sizes and applying thorough data cleaning and bias correction methods, as we did here.

Further refinement of our understanding of bee diversity across the state will require additional data collection or the application of novel modeling methodologies. Specifically, targeted surveys for rare bee species could enable the inclusion of additional bees of conservation interest in future models and facilitate assessments of the species that may be at the greatest risk of decline. The predicted distributions we present here provide an initial hypothesis of bee diversity and distributions across the state. These data products can aid in the protection of rare species by expediting the identification of new populations and streamlining targeted surveys of unmodeled species by prioritizing sampling in areas of high suitability for related modeled species or those sharing similar habitat and host plant preferences (Buckner & Danforth, 2022). Alternatively, with faster implementations of joint species distribution models on the horizon (Rahman et al., 2024), it may be possible to more accurately model rare species by borrowing strength from more common bees (Norberg et al., 2019). At a minimum, additional data collection through state surveys, national monitoring programs, or community science engagement will facilitate improved mapping of bee diversity, conservation, and ecosystem services, even if the resulting data is biased (Gaul et al., 2020).

The products resulting from our study represent the current best estimate of bee species richness and conservation habitat across NYS to date. By leveraging species distribution modeling methods robust to small sample sizes, we were able to model over half of NYS’s bee species. We present our results as gridded data layers to support statewide biodiversity conservation efforts. Our work serves as an example of the extensibility of statewide pollinator surveys and a roadmap for leveraging existing occurrence datasets to ensure that bee diversity is accounted for in contemporary biodiversity conservation initiatives.

## Acknowledgments

We thank Leif Richardson and Christopher Wilson for providing feedback on earlier iterations of our models. Members of the Pollinator Reading Group at Cornell University and the Danforth Lab shared valuable feedback on this manuscript. The Albany Pine Bush Preserve Commission, Katherine (Kass) Urban-Mead, and Leif Richardson provided additional bee occurrence records.

## Data Availability Statement

The raw occurrence data from the Empire State Native Pollinator Survey available from the New York Natural Heritage Program, restrictions on the use of this data may apply. A post-processing presence/background dataset is provided with permission of contributors for model reproducibility. The occurrence dataset and all data products produced during this study are openly available as GeoTIFFs stored on Figshare: https://doi.org/10.6084/m9.figshare.25883932 (Buckner et al., 2024a). All associated code is archived on Figshare: https://doi.org/10.6084/m9.figshare.25884169 (Buckner et al., 2024b).

