## Supplemental files for "Fine-Scale Models of Bee Species Diversity and Habitat in New York State"

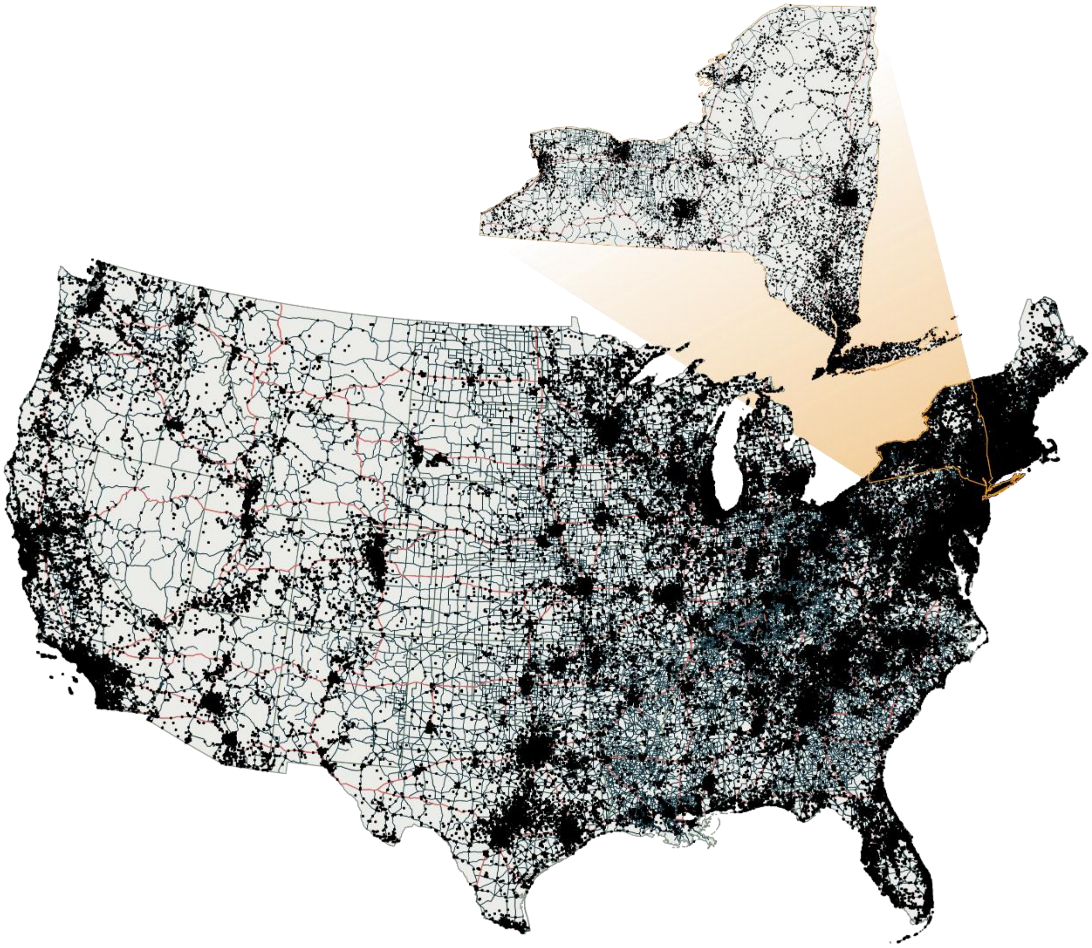

**Figure S1.** Distribution of presence records for all 269 bee species modeled across CONUS and within NYS. Primary and secondary roadways are shown as red and blue lines respectively.

**Table C1:** List of environmental covariates included in species distribution models for bee species across NYS before dimensionality reduction with a Principal Components Analysis (PCA). Covariates excluded from the PCA and models are marked with an "✗". Sampling Intensity was excluded from the PCA but was included in the models. All covariates used in the sampling intensity point process model are listed at the bottom of the table.

| Abbr. | Variable | Temporal Extent | Source | Excluded | Justification |
|---|---|---|---|---|---|
| *bio1* | Mean annual air temperature | 1981-2010 | [1,2] | | Warming temperatures may cause thermal stress, higher overwintering metabolic demands [3–5] |
| *bio7* | Annual range of air temperature | 1981-2010 | [1,2] | | Thermal variability across all life stages |
| *bio11* | Mean daily minimum air temperature of the coldest quarter | 1981-2010 | [1,2] | | Warming temperatures may cause higher overwintering metabolic demands, previously found to be important for a bee in region[3,6] |
| *bio12* | Annual precipitation amount | 1981-2010 | [1,2] | | Linked to variation in bee abundances[7] |
| *bio15* | Precipitation seasonality | 1981-2010 | [1,2] | | Within locality variability in precipitation may influence niche breadth[8] |
| *ai* | Aridity index | 1981-2010 | [1,2] | | Correlated with species richness in global analyses[9] |
| *ngd10* | Number of growing degree days above 10°C | 1981-2010 | [1,2] | | Approximates flight period/summer duration, may affect pre-wintering development and winter mortality[10] |
| *gsp* | Accumulated precipitation on growing season days | 1981-2010 | [1,2] | | Represents precipitation during the flight period, strong predictor in other analyses[7] |
| *lc* | Land cover (existing vegetation type) | 2014 | [11] | ✗ | Excluded from model, used only for preprocessing other covariates |
| *slp* | Slope | 2020 | [12] | ✗ | Excluded from model, used to calculate terrain roughness |
| *ssc* | Soil sand content | 1950-2017 | [13] | | Nest site selection in ground nesting species, may |

| Abbr. | Variable | Temporal Extent | Source | Excluded | Justification |
|---|---|---|---|---|---|
| | | | | | also serve as loose proxy for vegetation community[14] |
| *scc* | Soil clay content | 1950-2017 | [15] | | Nest site selection in ground nesting species, may also serve as loose proxy for vegetation community[14] |
| *rgh* | Terrain roughness (St. dev. of slope) | 2020 | [12] | | Proxy for microclimate availability and habitat heterogeneity[16] |
| *cc* | Canopy cover | 2014 | [17] | | Provide variable nest sites, floral resources, and distinct bee communities[18] |
| *int* | Predicted sampling intensity (point process model) | - | - | ~ | Model based bias correction |
| Sampling intensity model covariates | | | | | |
| *pop* | Population density | 2000 | [19] | | |
| *pad* | Protected areas | 2005-2022 | [20] | | |
| *mil* | Fort Drum boundary | 2010 | [21] | | |
| *road* | Distance to primary or secondary roadways | 2013 | [22] | | |
| *inst* | Distance to biodiversity institutions | - | [23] | | |
| *x* | Longitude | - | - | | |
| *y* | Latitude | - | - | | |

**Table C1 References:**


1. Karger, D. N. *et al.* Climatologies at high resolution for the earth's land surface areas. *Scientific Data* **4**, (2017).
2. Karger, D. N. *et al.* Climatologies at high resolution for the earth's land surface areas. EnviDat http://dx.doi.org/10.16904/envidat.228 (2021).
3. Williams, C. M., Henry, H. A. L. & Sinclair, B. J. Cold truths: how winter drives responses of terrestrial organisms to climate change. *Biological Reviews* **90**, 214–235 (2015).
4. Johnson, M. G., Glass, J. R., Dillon, M. E. & Harrison, J. F. How will climatic warming affect insect pollinators? in *Advances in Insect Physiology* vol. 64 1–115 (Elsevier, 2023).
5. Pardee, G. L. *et al.* Life-history traits predict responses of wild bees to climate variation. *Proceedings of the Royal Society B: Biological Sciences* **289**, (2022).
6. Buckner, M. A. & Danforth, B. N. Climate-driven range shifts of a rare specialist bee, Macropis nuda (Melittidae), and its host plant, Lysimachia ciliata (Primulaceae). *Global Ecology and Conservation* **37**, e02180 (2022).
7. Kammerer, M., Goslee, S. C., Douglas, M. R., Tooker, J. F. & Grozinger, C. M. Wild bees as winners and losers: Relative impacts of landscape composition, quality, and climate. *Global Change Biology* **27**, 1250–1265 (2021).
8. Quintero, I. & Wiens, J. J. What determines the climatic niche width of species? The role of spatial and temporal climatic variation in three vertebrate clades. *Global Ecology and Biogeography* **22**, 422–432 (2013).
9. Orr, M. C. *et al.* Global Patterns and Drivers of Bee Distribution. *Current Biology* **31**, 451-458.e4 (2021).
10. Sgolastra, F. *et al.* The long summer: Pre-wintering temperatures affect metabolic expenditure and winter survival in a solitary bee. *Journal of Insect Physiology* **57**, 1651–1659 (2011).
11. LANDFIRE. Existing Vegetation Type Layer. U.S. Department of the Interior, Geological Survey, and U.S. Department of Agriculture (2014).
12. LANDFIRE. Slope Degrees. U.S. Department of the Interior, Geological Survey, and U.S. Department of Agriculture (2020).
13. Hengl, T. Sand content in % (kg / kg) at 6 standard depths (0, 10, 30, 60, 100 and 200 cm) at 250 m resolution. Zenodo https://doi.org/10.5281/zenodo.2525662 (2018).
14. Antoine, C. M. & Forrest, J. R. K. Nesting habitat of ground-nesting bees: a review. *Ecological Entomology* **46**, 143–159 (2021).
15. Hengl, T. Clay content in % (kg / kg) at 6 standard depths (0, 10, 30, 60, 100 and 200 cm) at 250 m resolution. Zenodo https://doi.org/10.5281/ZENODO.2525663 (2018).
16. Title, P. O. & Bemmels, J. B. ENVIREM: an expanded set of bioclimatic and topographic variables increases flexibility and improves performance of ecological niche modeling. *Ecography* **41**, 291–307 (2018).
17. LANDFIRE. Forest Canopy Cover. U.S. Department of the Interior, Geological Survey, and U.S. Department of Agriculture (2014).
18. Ulyshen, M., Urban-Mead, K. R., Dorey, J. B. & Rivers, J. W. Forests are critically important to global pollinator diversity and enhance pollination in adjacent crops. *Biological Reviews* **98**, 1118–1141 (2023).
19. Center for International Earth Science Information Network - CIESIN - Columbia University. Gridded Population of the World, Version 4 (GPWv4): Population Density, Revision 11. NASA Socioeconomic Data and Applications Center (SEDAC) (2018).
20. USGS GAP. Protected Areas Database of the United States (PAD-US). U.S. Geological Survey data release https://doi.org/10.5066/P9Q9LQ4B (2022).
21. U.S. Census Bureau. 2010 TIGER/Line Shapefiles. (2010).
22. U.S. Census Bureau. 2013 TIGER/Line Shapefiles. (2013).
23. Zizka, A. *et al.* CoordinateCleaner: Standardized cleaning of occurrence records from biological collection databases. *Methods in Ecology and Evolution* **10**, 744–751 (2019).

**Table C2:** Principal Components Analysis loadings and summary statistics for each of the four principal components which were included in the species distribution models.

| Variable | PC1 | PC2 | PC3 | PC4 |
|---|---|---|---|---|
| *bio1* | 0.403 | -0.319 | 0.010 | -0.013 |
| *bio11* | 0.434 | -0.265 | -0.070 | 0.070 |
| *bio12* | 0.337 | 0.344 | 0.238 | 0.080 |
| *bio15* | -0.198 | -0.242 | -0.198 | 0.418 |
| *bio7* | -0.449 | 0.017 | 0.075 | -0.167 |
| *gsp* | 0.008 | 0.104 | -0.031 | 0.806 |
| *cc* | 0.236 | 0.416 | -0.007 | -0.133 |
| *ngd10* | 0.406 | -0.314 | -0.002 | -0.019 |
| *ai* | 0.232 | 0.445 | 0.184 | 0.121 |
| *scc* | -0.095 | -0.321 | 0.526 | 0.196 |
| *ssc* | 0.087 | -0.004 | -0.732 | -0.047 |
| *rgh* | -0.068 | 0.260 | -0.217 | 0.252 |

| Summary | PC1 | PC2 | PC3 | PC4 |
|---|---|---|---|---|
| Standard deviation | 1.984 | 1.754 | 1.255 | 1.041 |
| Proportion of Variance | 0.328 | 0.256 | 0.131 | 0.090 |
| Cumulative Proportion | 0.328 | 0.584 | 0.715 | 0.806 |

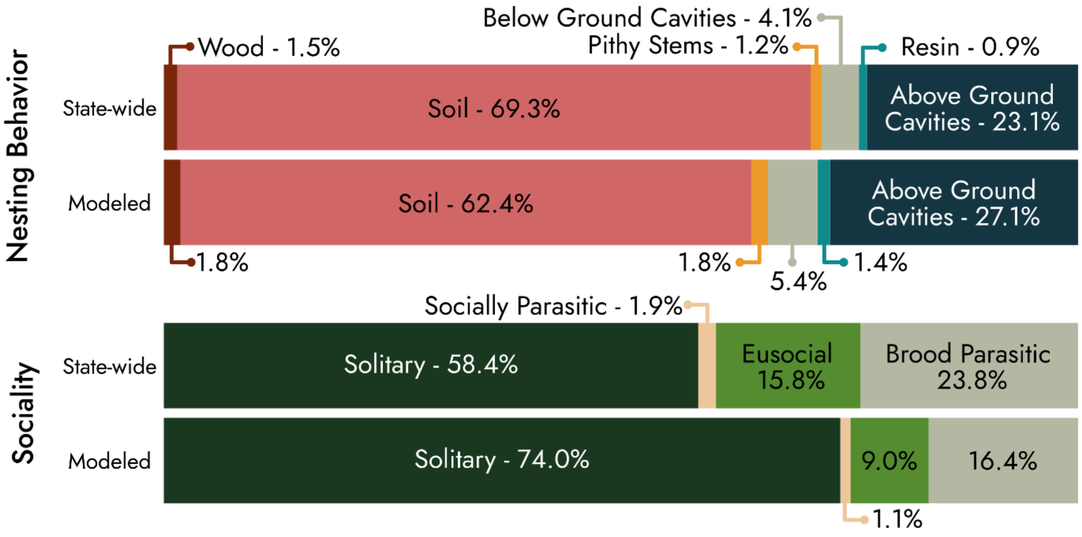


**Figure S2:** The percentage of species known from New York State (statewide) that display different nesting and social behaviors compared to the subset of species that were included in the models (modeled).

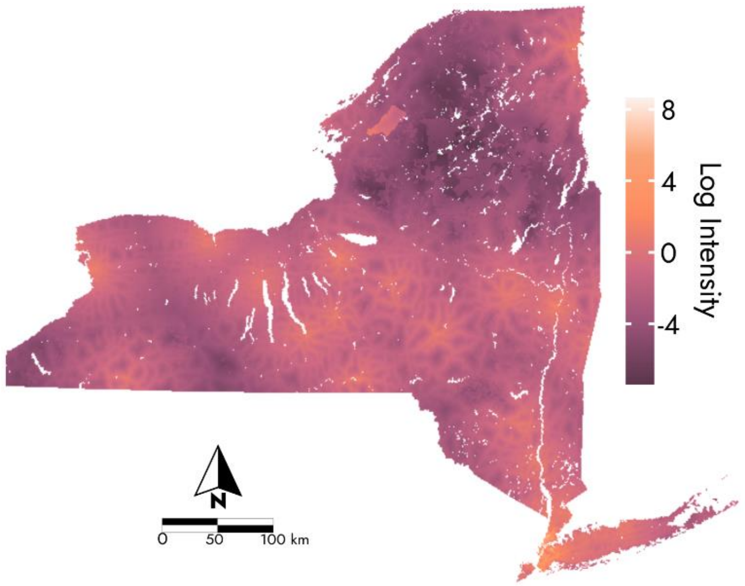


**Figure S3:** Log predicted sampling intensity across New York State showing high sampling intensity in New York City, Fort Drum, and along roadways near biodiversity institutions and the lowest values in remote areas of the Adirondack Mountains.

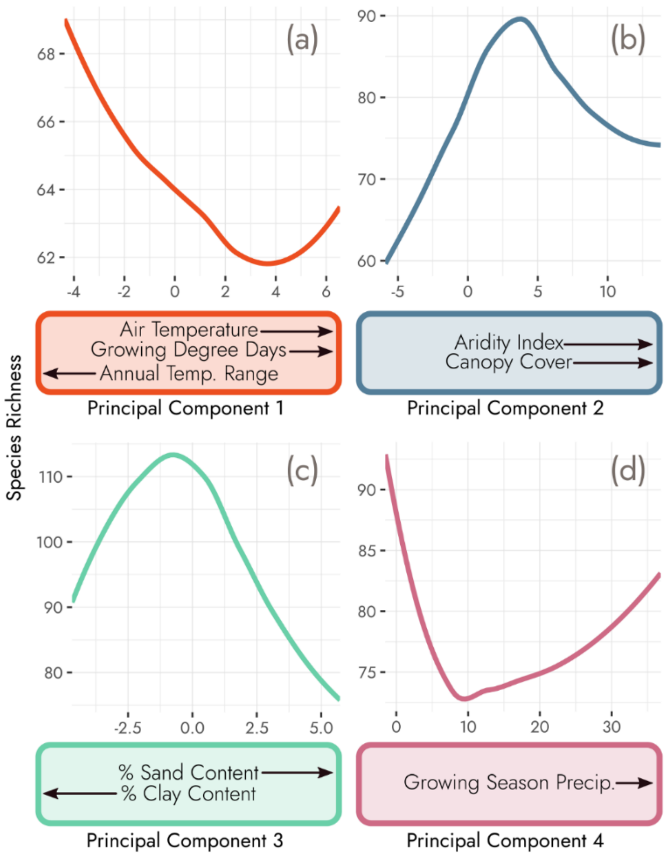


**Figure S4:** Species richness response curves for each of the four principal components (PCs). The environmental and climatic variables that contributed most to each PC and the direction of the relationship are included in the colored boxes. (a) PC1: species richness declines at higher temperatures (annual and winter) and decreasing variability. (b) PC2: Moderate aridity indices (higher moisture) and canopy cover support the highest predicted richness. (c) PC3: Richness peaks at intermediate soil textures. (d) PC4: Richness forms a U-shaped trend across growing season precipitation, with the greatest predicted richness when growing season precipitation is lowest.
